# Temporal changes in platelet activation and organ dysfunction during invasive *Streptococcus pyogenes* infection

**DOI:** 10.64898/2026.09.18.752656

**Authors:** Eleni Bratanis, Christofer Karlsson, Yashuan Chao, Zdenka Prgomet, Frida Palm, Johan Malmström, Oonagh Shannon

## Abstract

A key feature of sepsis is immune dysregulation caused when a local infection progresses to systemic disease. Innate immune cells, including platelets, can both contribute to immune defense but also to collateral organ damage. In this study, we characterized the molecular, cellular, and proteome responses at distinct stages of invasive *Streptococcus pyogenes* infection in a mouse model, from a local infection to dissemination and systemic inflammation. Invasion of bacteria to the bloodstream was associated with escalating cytokine levels, platelet activation, and an early leukocytosis that progressed to leukopenia. Multiparameter mapping of organ function using plasma biomarkers, histopathological changes, and proteome reorganization revealed distinct temporal patterns of organ dysfunction. Platelets were mobilized to the local skin infection at early stages of inflammation but also accumulated in the liver at later stages, coinciding with sepsis-induced organ damage. We demonstrated that *S. pyogenes* bound to the GPIbα receptor on platelets *in vitro*, suggesting that bacteria may bind to platelets in the bloodstream. Treatment of infected mice with antibiotics diminished the bacterial load, reduced sepsis-induced organ damage, and partially reverted the disease-associated proteome rearrangement in the plasma and liver. Treatment of infected mice with pharmacological blockade of platelet activation with ticagrelor neither exacerbated disease nor significantly protected against sepsis-induced organ damage. However, at the proteome level, ticagrelor significantly reverted sepsis-induced platelet activation and stress responses in the liver. We conclude that platelets contribute to liver dysfunction in sepsis and may represent a target for adjunct therapy in combination with antibiotics.

## Introduction

Sepsis, a life-threatening organ dysfunction caused by a dysregulated host response to infection ^1^ is a leading cause of critical illness and mortality worldwide ^2^. Central to sepsis pathophysiology is an excessive systemic inflammatory response (SIRS), followed by a compensatory anti-inflammatory response (CARS) and organ dysfunction ^1,3,4^. The disease progression is, therefore, complex and involves dysregulation of multiple host-pathogen interaction interfaces at the local site of infection, in the bloodstream, and in distant organs. *Streptococcus pyogenes* is a significant human pathogen that causes both mild skin infections but also severe invasive disease and sepsis ^19^. Current management of sepsis includes prompt antimicrobial therapy and a limited use of broad-spectrum immunomodulatory therapies. Enhanced understanding of sepsis pathophysiology in recent years has not yet translated into improved treatments ^5^. A contributing factor is the poor translational power of mouse models of sepsis where mainly fulminant models of uncontrolled inflammation are generated by systemic administration of bacteria or toxins. Such models do not adequately reflect the transition from a local controlled inflammation to sepsis.

Local inflammation involves a coordinated recruitment of immune cells and mediators from the blood into the infected tissue. However, if this local response fails to limit infection, bacteria can gain access to the bloodstream. The ensuing systemic inflammation results in endothelial damage and activation of circulating leukocytes, the coagulation system, and platelets. A hallmark of sepsis is dysregulation of coagulation and inflammation, which can ultimately result in disseminated intravascular coagulation (DIC), characterized by microvascular thrombosis and bleeding ^10,11^. Platelets have a fundamental role in both coagulation and immune responses. Platelets respond directly to invading pathogens through surface receptors, including Toll-like receptors (TLRs), complement receptors, and Fc receptors ^6, 7, 8^. Multifaceted roles have been attributed to platelets in sepsis ^9^; therefore, platelet activation is a pharmacological target of interest in the treatment of sepsis ^16^. However, the therapeutic value of antiplatelet agents in sepsis needs to be assessed in relation to an associated risk of increased bleeding. Thrombocytopenia occurs in sepsis, is an independent risk factor for mortality, and is incorporated into the Sequential Organ Failure Assessment (SOFA) score ^12,13^. The events preceding thrombocytopenia and the role of platelets in local immune defense have, however, not been clarified *in vivo*. For example, platelet depletion in a sepsis model following lung infection with *Klebsiella pneumoniae* resulted in increased bacterial load and mortality ^14^. Conversely, platelet depletion prior to intraperitoneal infection with *S. pyogenes* was associated with decreased bacterial survival and dissemination ^15^. Together, these findings suggest that platelet responses are dynamic during infection and warrant further investigation during the progression from local infection to sepsis.

In the present study, we performed multiparameter mapping of the host response in a mouse model of invasive *S. pyogenes* infection during the progression from a local skin infection to a systemic inflammatory syndrome and organ dysfunction, with a particular focus on platelet responses during infection and sepsis. Time-resolved molecular signatures reflecting immunological, coagulation, and organ dysfunction were mapped in blood and organs using quantitative mass spectrometry-based proteomics, flow cytometry, and immunohistochemistry. These approaches were further applied to evaluate the impact of therapeutic interventions on disease progression by comparing gold-standard antibiotic therapy using meropenem with antiplatelet therapy using the platelet P2Y_12_ receptor antagonist ticagrelor.

## Methods

### Bacteria and culture conditions

*Streptococcus pyogenes* AP1 (from the Collection of the World Health Organization Collaborating Center for Reference and Research on Streptococci, Prague, Czech Republic) were grown in Todd– Hewitt broth, supplemented with 0.2 % yeast extract (THY, BD diagnostic), overnight at 37°C and 5% CO_2_.

### Streptococcus pyogenes infection in mice

All animal use and procedures were approved by the local Malmö/Lund Institutional Animal Care and Use Committee, ethical permit number 03681-2019. *S. pyogenes* AP1 was grown statically to logarithmic phase in THY medium (37°C, 5% CO_2_). Bacteria were washed and resuspended in sterile PBS. Nine-week-old female C57BL/6J mice (Janvier, Le Genest-Saint-Isle, France) were shaved and infected with 50 µl (containing 2 x 10^5^ cfu) bacteria by subcutaneous injection on the right flank. The control group was similarly injected with sterile PBS (n=9). Mice were rehydrated subcutaneously in the scruff with saline at 24 h post-infection (hpi). Body weight and general symptoms of infection were monitored regularly. Groups of mice were sacrificed at 6 hpi (n=10), 12 hpi (n=10), 24 hpi (n=10), and 36 hpi (n=15), and organs (blood, skin, liver, spleen, and kidney) were collected. Blood was taken by cardiac puncture and collected in tubes containing sodium citrate (MiniCollect tube, Greiner Bio-One). For treatment interventions, mice received either the antibacterial meropenem (10 mg/kg) at 12 hpi or the antiplatelet ticagrelor (30 mg/kg) at 12 and 24 hpi by intraperitoneal injection. Treated mice were sacrificed at 36 hpi, and organs (blood, skin, liver, spleen, and kidney) organs were collected.

### Organ preparations

Citrated blood collected from infected and control mice was centrifuged (2000 rcf, 10 min) to obtain platelet free plasma. Plasma was aliquoted and stored at -80°C. Collected organs were homogenized (MagnaLyzer, Roche) in PBS using sterile silica beads (1 mm diameter, Techtum). Skin biopsies (10 mm, Acu punch) and spleens were either prepared for histology or homogenized for plating and proteomics. Independent liver biopsies and kidney biopsies were prepared in parallel for both histology and homogenization. Degree of bacterial dissemination was determined by serial diluting and plating (skin, liver, kidney and spleen homogenates) onto blood agar plates. Colony forming units were counted following overnight incubation (37°C, 5% CO_2_) and are presented as cfu/g of tissue. Remaining organ homogenates were centrifuged (20000 rcf, 10 min) and supernatants immediately transferred and aliquoted into fresh vials. All samples were stored at −80 °C until further analysis. Protein concentration was determined using standard BCA assay (Thermo Scientific) according to manufacturer instructions.

### Flow cytometry of blood cells

Citrated blood collected from infected and control mice was diluted with HEPES buffer containing Mouse BD Fc-block (BD Pharmingen). Platelets, monocytes and neutrophils were identified using, anti-CD45 FITC (BD Pharmingen), anti-CD41 FITC (BD Pharmingen), anti-Ly6G PE (BD Pharmingen), anti-Ly6C APC-R700 (BD Horizon) and anti-CD11b PerCP (BD Pharmingen). All antibodies were diluted 1:200 and incubated (15 min, RT). Samples were lysed using 1-step Fix/Lyse Solution (e-Bioscience), washed (500 rcf, 5 min) cellular pellets were resuspended in PBS. The samples were analyzed using an Accuri Plus C6 Flow Cytometer (BD Biosciences), and the data was analyzed using C6 Software (BD Biosciences). Total leukocytes were gated according to characteristic forward and side scatter. Neutrophils were identified as CD11b high, Ly6G high and Lyg6C low. Monocytes were identified as CD11b high, Ly6G negative and Lyg6C intermediate/high. The gating strategy is shown in Supplementary Figure 4).

Platelets and platelet activation following treatment with Mem or ticagrelor were separately analyzed following activation with thrombin (1U), using anti-CD62P FITC (Emfret ANALYTICS), anti-Jon/A PE (Emfret ANALYTICS) and anti-CD41 APC (BD Pharmingen) antibodies.

### Plasma cytokines and organ damage markers

Cytokine levels in citrated plasma samples were quantified using a cytometric bead array assay (CBA mouse inflammation kit, BD) according to manufacturer instructions and analyzed using a FACSVerse (BD Biosciences). Mouse plasma levels of alanine aminotransferease (ALT), lactate dehydrogenase (LDH), blood urea nitrogen (BUN), hyaluronic acid (HA) and soluble P-selectin/ CD62 were measured using Alanine Transaminase Activity Assay Kit (ab105134, Abcam), Lactate Dehydrogenase (LDH) Assay Kit (ab102526, Abcam), Urea Nitrogen (BUN) Colorimetric Detection Kit (EIABUN, Thermo Fischer Scientific), Hyaluronan Quantikine ELISA Kit (DHYAL0, R&D systems) and Mouse P-selectin/CD62 (DY737, R&D systems), respectively. All ELISA and colorimetric assays were performed according to the manufacturer instructions in combination with a microplate reader (Victor 3 Multilabel Plate Reader, Perkin Elmer).

### Pathology scoring of organs

Skin, liver, and kidney tissue samples from infected and control mice were fixed in Histofix (Histolab Products AB, Askim, Sweden) for 48 h, dehydrated in 70% EtOH for at least 24 h, embedded into paraffin blocks and sectioned (4 μM microtome, Leica RM2255). Sections were transferred to slides and deparaffinized by incubation (60°C, 30 min) followed by dehydration in Histolab clear (#14250, Histolab). Sections were stained using Mayer hematoxylin and eosin (H&E) (Histolab Products AB, Askim, Sweden). Sections were air-dried at RT and mounted with Pertex 5 (#00840, Histolab) and cover glass (1.5 mm). Imaging was performed using a light microscope (Nikon Eclipse 80i: 10x magnification in skin and 40x magnification in kidney and liver).

Representative images from haematoxylin and eosin-stained sections of liver and kidneys (n=3 per time point) were acquired at 20x and 40x magnifications. Image analysis of the sections was performed with FIJI version 1.53. For quantification of liver dysfunction, the presence of clots in the liver vessels was counted manually. For quantification of kidney dysfunction, mesangial proliferation in kidneys was assessed. A minimum of 36 glomeruli were selected using the freehand tool and processed with Huang chosen as the method for auto threshold (T0, n= 45; T6, n=36; T12, n=43; T24, n=41; and T36, n=39). Images were deconvolved using the colour deconvolution function, with vectors set to H&E. The image depicting haematoxylin staining was selected and auto threshold set to Otsu was applied. The threshold image was added to the ROI manager and then objects within the ROI were measured. Representative images from haematoxylin and eosin-stained sections of skin biopsies were acquired at 4x and 10x magnifications.

### Immunohistochemistry of platelets in tissue

Formalin-fixed paraffin-embedded mouse skin and liver tissues (n=5 per time point) were consecutively sectioned at 3 μm and at first stained with hematoxylin-eosin (H&E) according to standard protocol.

To detect CD41 expression, tissue sections were deparaffinized and rehydrated prior to heat-induced antigen retrieval in TRIS-EDTA buffer, pH 9.0 at 95°C for 20 min in a Decloaking Chamber^TM^ (NxGen, Biocare Medical). Non-specific background staining was blocked with Background Punisher (MACH4, BRI4012L, Biocare Medical) prior to overnight incubation at 4°C with CD41 antibody (clone EPR4330, ab134131, Abcam) diluted at 1/500 with antibody diluent (S2022; DAKO). Next, endogenous peroxidase was blocked with peroxidase blocking reagent (S2023; DAKO) prior to incubation with secondary antibody (goat anti-rabbit-HRP, MACH4). Immunoreaction was visualized with diaminobenzidine (DAB, MACH4). Tissue sections were then counterstained with hematoxylin, dehydrated, and mounted. After each step, tissue sections were washed with Tris-buffered saline, 0.1 % Tween 20, pH 7.6. Tonsil was used as a positive control for CD41 expression, while N-Universal rabbit negative control (N1699; DAKO) and omission of the primary antibody served as negative controls. Stained tissue sections were scanned and analyzed using Nano Zoomer histology scanner (Hamamatsu) using ×40 objective.

### *In vitro* platelet-bacteria complex formation

Citrated blood collected from infected and control mice were diluted with HEPES buffer containing Mouse BD Fc-block (BD Pharmingen). *S. pyogenes* AP1 was grown to logarithmic phase in THY medium (37°C, 5% CO_2_). Bacteria were washed, resuspended in sterile PBS, and adjusted to 2 x 10^7^ cfu/ml. Samples were incubated with the platelet receptor-blocking antibodies anti-GPIbα (Emfret) or anti-GPIIb/IIIa (Emfret), the GPIIb/IIIa antagonist Integrilin (100 μg/ml), or the P2Y_12_ antagonist ticagrelor (Brilique; 0.5 mg/ml). Isotype control antibodies, rat IgG2b and rat IgG2a (Miltenyi Biotec) were also included. Platelet-bacteria complex formation was identified by anti-CD41 APC (BD Biosciences) and anti-*S. pyogenes* FITC (abcam). Antibodies were diluted (1:200 anti-CD41 and 1:50 anti-GAS) and incubated with the samples (15 min, RT). Samples were lysed using 1-step Fix/Lyse Solution (e-Bioscience) and washed (500 rcf, 5 min). Samples were resuspended in PBS and analyzed using an Accuri C6 Plus Flow Cytometer (BD Biosciences). The data was analyzed using C6 Software (BD Biosciences).

### MS sample preparation and data acquisition

Leukocytes (n = 24) were isolated from infected and control mice using the cell pellet remaining after plasma collection and the AllPrep Mini Kit (Qiagen). The protein-containing flow-through obtained after total RNA isolation was precipitated using three volumes of ice-cold acetone and incubated for 1 h at -18 °C. Precipitates were then centrifuged for 15 min at 14,000 *g*, 4 °C after which the supernatant was discarded, and the protein pellet was air-dried. The protein precipitates were subsequently dissolved and denatured in 50 µl 0.1% RapiGest SF (Waters), 8 M urea (Sigma-Aldrich) and 100 mM ammonium bicarbonate and followed the digestion protocol for the other sample types below. Plasma (n=46) and organ homogenates (liver n=48, kidney n=24 and skin n=47) were denatured with 8M urea and reduced with 5 mM Tris(2-carboxyethyl)phosphine hydrochloride, pH 7.0 for 45 min at 37 °C, and alkylated with 25 mM iodoacetamide (Sigma) for 30 min followed by dilution with 100 mM ammonium bicarbonate to a final urea concentration below 1.5 M. Proteins were digested by incubation with trypsin (1/100, w/w, Sequencing Grade Modified Trypsin, Porcine; Promega) for at least 9 h at 37 °C. Digestion was stopped using 5% trifluoracetic acid (Sigma) to pH 2 to 3. Peptide clean-up was performed by C18 reversed-phase spin columns according to manufacturer instructions (Silica 10 µg C18 300 Å Columns; Harvard Apparatus). Solvents were removed using a vacuum concentrator (Genevac, miVac) and samples were resuspended in 50 μl HPLC-water (Fisher Chemical) with 2% acetonitrile and 0.2% formic acid (Sigma). Peptide analyses (n=189) were performed on a Q Exactive HF-X mass spectrometer (Thermo Fisher Scientific) connected to an EASY-nLC 1200 ultra-HPLC system (Thermo Fisher Scientific). Peptides were trapped on precolumn (PepMap100 C18 3 μm; 75 μm × 2 cm; Thermo Fisher Scientific) and separated on an EASY-Spray column (ES903, column temperature 45 °C; Thermo Fisher Scientific). Equilibrations of columns and sample loading were performed per manufacturer’s guidelines. Mobile phases of solvent A (0.1% formic acid), and solvent B (0.1% formic acid, 80% acetonitrile) was used to run a linear gradient from 5% to 38% over 90 min at a flow rate of 350 nl/min. The data independent acquisition (DIA) method is described by Bruderer et al^40^. In summary, one MS1 scan with a scan range of 350–1650 m/z at resolution of 120,000 was followed by 44 DIA MS2 isolation scans with at resolution of 30,000 and with a stepped normalized collision energy (NCE) of 25.5, 27 and 30.

### DIA data analysis

Raw files were converted to mzML with ThermoRawFileParser v1.3.1 and searched with DIA-NN v1.8.1. A hybrid spectral library strategy was used. Where the existing HF-X experimental mouse library was from Mohanty et al. (2023) and was reannotated against the mouse reference proteome (UniProt UP000000589, release 2022_02; 21,953 protein sequences) and a deep-learning-predicted in silico library was generated from the same FASTA. DIA-NN then re-analysed the runs against this newly generated library (‘--reanalyse’), retaining predicted spectra where they were judged more reliable than experimental ones.

The data was searched batch-wise with parameters: trypsin specificity with up to one missed cleavage and N-terminal methionine excision; carbamidomethylation of cysteine as a fixed modification and no variable modifications; peptide length 7–30 residues; precursor *m/z* 350–1650; precursor charge 2–4; fragment *m/z* 200–1800. Mass accuracy was determined automatically by DIA-NN per batch. Precursor and protein-group identifications were filtered to 1% FDR and protein quantities were derived by the MaxLFQ algorithm with cross-run normalisation.

### Bioinformatics analysis

MS data analysis was performed using custom scripts in R (4.1.12). The DIA-NN protein and precursor matrix reports were used for downstream analysis. Protein quantifications were filtered, requiring >75 % id rates in at least one sample group per batch. Missing protein quantification values was assumed to be left-censored were imputed using the Quantile Regression Imputation of Left-Censored data method with imputeLCMD v2.1 R package. Differential abundance testing was performed with DEqMS v1.11.1 R package using the filtered protein intensities and peptide per protein counts from DIA-NN matrices. Cut-offs were log2 foldchange > ±1.5 and adjusted P-value < 0.01 for infection response and adjusted P-value < 0.05 for intervention response. Data was visualized with t-distributed Stochastic Neighbor Embedding (t-SNE) method using the Rtsne R v 0.16 package. Heatmaps were generated with ComplexHeatmap R package v 2.10 using Ward.D2 clustering method. Treatment efficacy was assessed using a proteome reversion framework adapted from Mohanty et al. (2023), in which infection-associated protein abundance changes (healthy vs. untreated disease) were regressed against treatment-induced changes (untreated vs. treated). Functional and pathway enrichment analysis was performed with Metascape^23^ using the web interface (https://metascape.org/) and “Express Analysis of Multiple Gene Lists” workflow.

### Statistical analysis

Statistical analysis was, unless otherwise stated, performed using nonparametric Mann-Whitney tests (Prism 9.1.0 software; GraphPad, Inc). P values < 0.05 were considered statistically significant.

## Results

### Mobilization of the innate immune response during disease progression

Subcutaneous skin infection was induced in mice, and disease progression was monitored over time by sacrificing groups at defined endpoints (Fig. 1A). Increasing weight loss was observed at each endpoint, reaching a mean of 10% decrease already at 24 hours post-infection (hpi) (Fig. 1B). Following an initial increase, the local bacterial burden in the skin plateaued at 12 hpi. At the same time point, significant bacterial dissemination was evident to the liver, spleen, and kidneys and continued to increase over time (Fig. 1C). In plasma, significant increases in pro-inflammatory mediators were detected at 12 hpi (Fig. 1D). Plasma IL-6 and TNF-α levels continued to increase throughout the later stages of infection, indicating a progressive systemic inflammatory response, whereas MCP-1 and IFN-γ peaked at 12 and 24 hpi, respectively. The total number of circulating leukocytes in blood rapidly increased in response to skin infection, accompanied by proportional increases in neutrophils and monocytes (Fig. 1E). As systemic infection progressed, leukopenia, in particular neutropenia, developed at 24 hpi (Fig. 1E). Platelet activation and degranulation occurred late during infection (24 hpi), as indicated by increased plasma CD62P levels (Fig. 1E). Collectively, these findings demonstrate a progressively dysregulated immune response characterized by excessive cytokine/chemokine levels, platelet activation, and leukopenia—a hallmark of sepsis.

**Figure 1.**
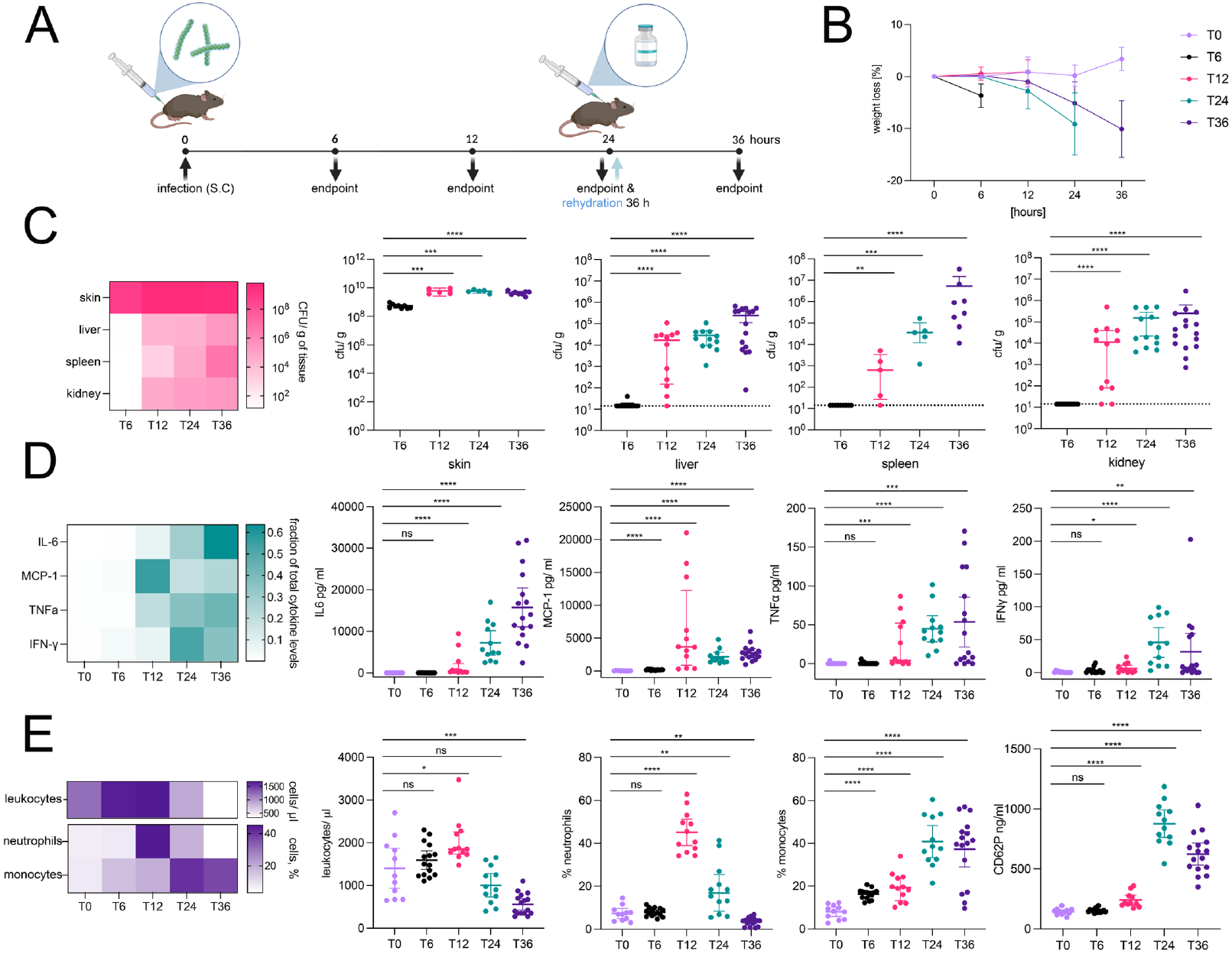
*S. pyogenes* skin infection results in increased bacterial dissemination, pro-inflammatory cytokine levels, and blood cell mobilization. (A) Nine-week-old female C57BL/6J mice were infected subcutaneously on the flank with *S. pyogenes* AP1 or PBS and rehydrated at 24 hours post-infection (hpi). Animals were sacrificed at 6, 12, 24, or 36 hpi. Blood, skin, liver, spleen, and kidney were harvested at each time point. (B) Weight loss was determined at each time point 6, 12, 24, or 36 hpi. T0 weights represents healthy controls at each time point (C) Bacterial load (cfu/g) was determined by serial dilution and viable count determination after overnight incubation. (D) Cytokine levels (pg/ml) in plasma were measured using a cytometric bead array. (E) Blood cell distribution was analyzed by flow cytometry to determine total leukocyte counts (cells/µl) and the relative proportions of neutrophils and monocytes (% of the leukocytes). Heatmaps display mean cfu/g of tissue (C), mean fraction of total cytokine levels in plasma (D), and mean total leukocyte counts (cells/ µl) and percentage of neutrophils and monocytes (E). Cfu/g of tissue in skin and spleen were determined at T6 (n=9), T12 (n=5), T24 (n=5), and T36 (n=8), and in liver and kidney at T6 (n=15), T12 (n=12), T24 (n=12), and T36 (n=16). Cytokine levels, blood cell distribution, and plasma CD62P levels were determined at T0 (healthy controls; n =11), T6 (n=15), T12 (n=12), T24 (n=12), and T36 (n=16). Dotted lines, if present, represent detection limits for the assays.

### Biomarkers of sepsis induced organ damage in plasma and tissue

The time-resolved impact of *S. pyogenes* infection at the local site and at distant organs over time was assessed by quantitative mass spectrometry-based proteomics. Data-independent acquisition mass spectrometry (DIA-MS) and targeted data extraction (SWATH-MS) demonstrated organ-specific dynamics in kidney, leukocytes, liver, plasma, and skin samples from infected mice (6, 12, 24, and 36 hpi) as compared to uninfected controls (0 hpi). Time-dependent proteome changes were observed across organs and plasma (Fig. 2A). The most pronounced temporal separation occurred between 12 and 24 hpi across all organs, although early separation was already evident in kidney, leukocytes, and skin. Separation from baseline (0 hpi; uninfected controls) was most evident during the later stages of infection (24 and 36 hpi). In contrast, the local site of infection, skin, showed substantial early proteome changes already at 6-12 hpi. Differential protein abundance analysis revealed a temporal increase in the number of differentially abundant proteins (DAPs), with increasing numbers of both up- and downregulated proteins over time (Fig. 2B). The full list of proteins is included as Supplementary Table 1. The blood leukocyte pool exhibited a predominance of downregulated proteins during later stages of infection, which likely reflects leukocyte activation and overall changes in the circulating pool during extravasation of leukocytes from blood into tissue. The liver exhibited the greatest number of DAPs, with a peak of approximately 900 upregulated proteins at 24 hpi, followed by approximately 800 downregulated proteins at 36 hpi. Among the upregulated proteins were acute-phase response proteins such as serum amyloid A, ferritin, mannose binding protein, complement factors, and fibrinogen. At 24 and 36 hpi, there was a downregulation of proteins associated with some metabolic pathways, such as lipid metabolism and mitochondrial function, and upregulation of proteins associated with cellular stress, including heat shock proteins and MAPK family proteins. Approximately 200 proteins were upregulated in plasma at 24 and 36 hpi including acute-phase reactants and immune mediators. Similarly, skin exhibited approximately 400 upregulated proteins at each time point from 12 hpi onwards, indicating an early and persistent response to infection in the tissue. Collectively, these findings demonstrate progressive organ-specific proteome remodeling.

**Figure 2.**
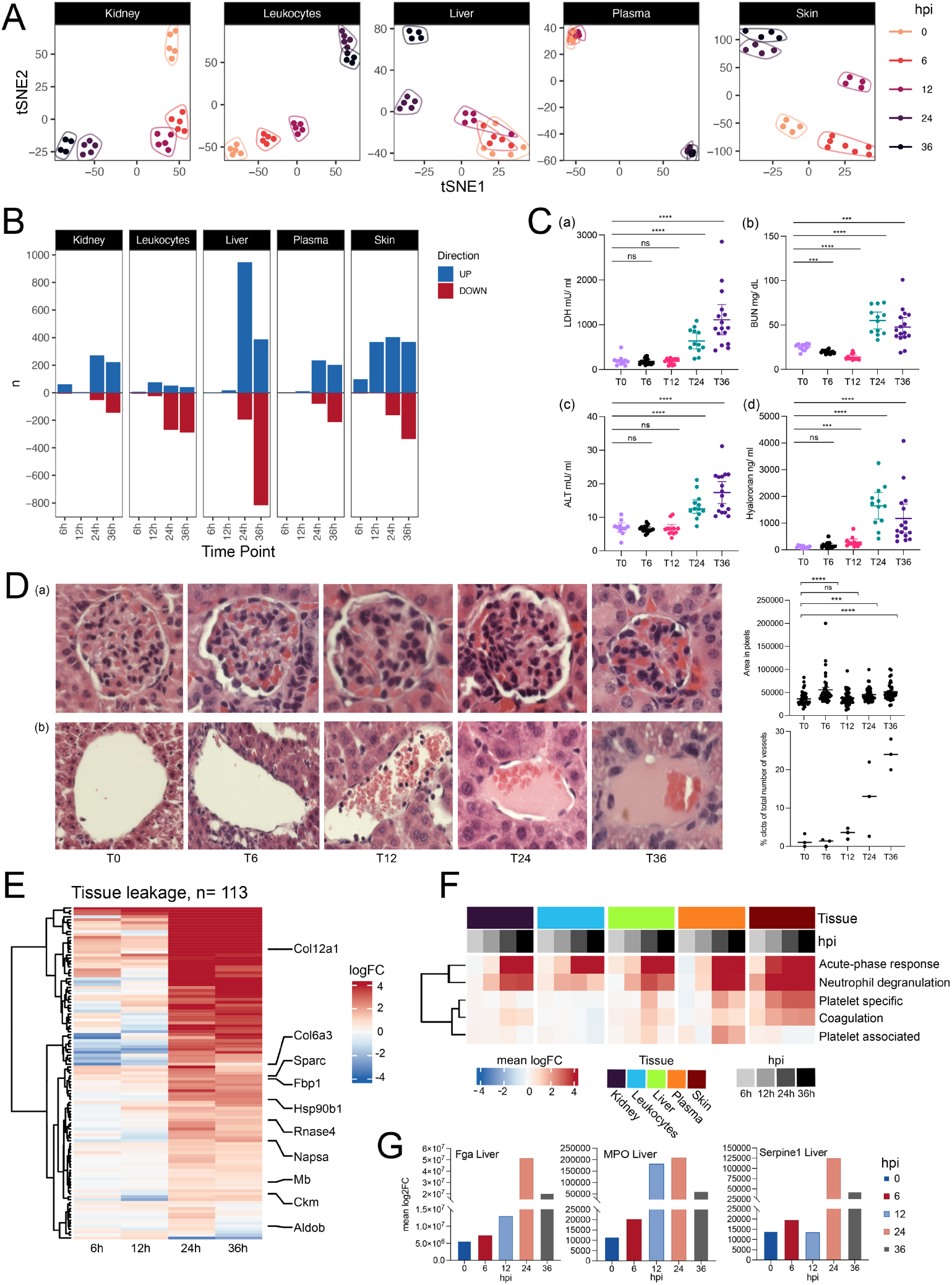
Local *S. pyogenes* infection progresses to sepsis-induced organ damage and tissue leakage. The kinetics of organ damage in infected mice was assessed using a combination of proteome profiling with direct pathological assessment of tissue and plasma biomarkers of organ damage. (A) tSNE plots of the full proteome. The time points are indicated by colors from 0 to 36 hours post-infection (hpi). (B) The number of differentially abundant proteins (DAPs) in the organs over time. (C) Plasma levels of organ damage markers (a) lactate dehydrogenase (LDH), (b) blood urea nitrogen (BUN), (c) alanine transaminase activity (ALT), and (d) hyaluronic acid (HA). (D) Pathological assessment of organ damage in H&E-stained biopsies. Representative images illustrate the qualitative changes over time in (a) kidney and (b) liver. Quantitative assessment of damage was determined as (a) increasing mesangial cell proliferation in the glomeruli and (b) hepatic intravascular coagulation (% of vessels with clots). Quantifications were performed using ImageJ (FIJI version 1.53) and are presented as (a) area in pixels and (b) % clots of total number of vessels. (E) Heatmap showing log2 fold changes compared to 0 hpi of 113 tissue-abundant and tissue-specific proteins in plasma samples over time. (F) The DAPs were subjected to functional enrichment using Metascape. The total log2 fold change for selected enrichments are displayed in a heatmap for kidney, leukocytes, liver, plasma, and skin for all time points. (G) Temporal changes in marker proteins associated with coagulation and neutrophil accumulation in liver tissue.

To assess sepsis-associated organ dysfunction, established markers of organ damage were measured in plasma (Fig. 2C). Lactate dehydrogenase (LDH), a general marker of cell and tissue injury, increased at 24 hpi and reached a peak at 36 hpi (Fig. 2Ca). Plasma levels of blood urea nitrogen (BUN) and alanine transaminase activity (ALT) are associated with renal and liver damage, respectively. Both markers were stable during the early host response and increased at 24 hpi (Fig. 2Cb-c). At 36 hpi, BUN levels decreased slightly while ALT levels increased. Endothelial cells are important drivers of microvascular dysfunction in organ damage. To assess endothelial injury, hyaluronic acid (HA) levels in plasma were determined, as HA has previously been reported to be elevated in septic mice (Fig. 2Cd) ^20^. Plasma HA levels increased already at 12 hpi, peaked at 24 hpi, and decreased slightly at the final time point (Fig. 2Cd). Organ pathology was characterized in hematoxylin and eosin (H&E)-stained tissue sections from kidney (Fig. 2Da) and liver (Fig. 2Db). Structural alterations were observed in kidney over time, including fibrin clot accumulation within the glomeruli. Quantitative histopathological assessment of renal damage based on mesangial cell proliferation in the glomeruli revealed increased mesangial cell proliferation at 24 hpi that increased further at 36 hpi (Fig. 2Da), consistent with ongoing inflammation and tissue damage. Morphological changes in the liver were already detected at 12 hpi, including leukocyte accumulation in the hepatic vessels, fibrin clot formation in vessels, and tissue necrosis. Quantitative histopathological assessment of liver damage based on manual counting of intravascular clot formation in 100 vessels per liver section revealed increased clot formation at 24 hpi (Fig. 2Db). Clot formation and thrombosis continued to increase in the liver over time; however, there was a heterogenous response within groups. Collectively, these findings demonstrate progressive organ dysfunction and tissue damage.

The kinetics of these traditional markers of organ damage corresponded to the pronounced proteomic reorganization of organ tissue observed at 24 and 36 hpi (Fig. 2A and 2B). To further assess protein biomarkers of sepsis-induced organ damage, differentially abundant tissue proteins in plasma were categorized using protein tissue origin inferred from a previously published murine/human atlas ^21,22^, thereby defining a tissue leakage signature (Fig. 2E). The plasma abundance of 113 tissue-associated proteins (Supplementary Table 2) increased over time, with predominant tissue leakage occurring at 24 hpi that plateaued at 36 hpi, including proteins from heart (Mb, Ckm), lungs (Sparc, Col6a3), liver (Hsp90b1, Fbp1, Aldob), and kidney (Napsa, Col12a, Rnase4, Aldob) (Fig. 2E). Critical molecular features of infection were further identified in the tissues and in plasma using functional enrichment analysis of DAPs, performed using Metascape ^23^ (Fig. 2F). The local site of infection, skin, showed an early accumulation of proteins (12 hpi) associated with the acute-phase response, neutrophil degranulation, coagulation, and platelets. This reflects a robust acute inflammatory response at the local site. The same classes of acute-phase reactants were not elevated in plasma until a later time point (24 hpi). At 24 and 36 hpi, proteins associated with the acute-phase response and neutrophil degranulation were the predominantly abundant classes observed in kidney, liver, and leukocytes, while coagulation and platelet proteins were less enriched in these organs (Fig. 2F). This may reflect the transition of the liver to producing acute-phase reactants required for the ongoing infection but also an infiltration of neutrophils into the organ. Proteins associated with coagulation, such as Fga, and neutrophil activation, such as MPO and Serpine 1, were most enriched in the liver at 24 hpi (Fig. 2F and 2G), which agreed with histopathological changes observed in the liver (Fig. 2D). Collectively, the combination of pathophysiological and proteome data maps the temporal changes from an initial local infection that progresses to organ damage and sepsis over time.

### Skin infection is associated with leukocyte, platelet, and plasma protein accumulation in tissue

Initial macroscopic changes observed at the local site following infection included cardinal signs of inflammation such as swelling and reddening of the skin. The inflammation-induced pathological changes at the local infection site were investigated using hematoxylin and eosin (H&E) staining of skin biopsies (Fig. 3A). Within 6-12 hpi, local disruption of the dermal layers was observed in the skin. Leukocyte infiltration from blood vessels into the tissue clearly increased over time, indicating vascular changes and leukocyte recruitment from the vasculature. Extensive necrosis was observed at 24-36 hpi, with clear disruption of the dermal layers. The presence of platelets at the local site of inflammation was investigated using immunohistochemistry (Fig. 3B). Platelets were present mainly as single cells in local vessels at 6 and 12 hpi, however, some aggregates of platelets accumulated in the vessels at 24 and 36 hpi, which coincided with increased tissue necrosis observed at the same time. Platelets remained confined to the vasculature (Fig. 3B), in contrast to the widespread neutrophil infiltration observed in H&E-staining of the same tissue (Fig. 3A). Global proteome changes in the skin at the same time points revealed an enrichment of proteins associated with neutrophil degranulation, such as elastase (Fig. 3Ca and Fig. 3D). An accumulation of histones, such as histone 4, at 12-36 hpi likely reflects cell necrosis in the tissue (Fig. 3D). This was accompanied by an enrichment of proteins involved in coagulation and acute-phase response over time (Fig. 3Cb-e). Furthermore, platelet proteins involved in surface receptor activation (GpIb, CD36), cytoskeleton rearrangement (Actn1, Plek), and granule release (Pf4, Thrombospondin) increased rapidly at 12-24 hpi (Fig. 3Ce and Fig. 3D). Collectively, these findings demonstrate progressive local tissue inflammation characterized by neutrophil infiltration, vascular platelet accumulation, and associated proteome changes during invasive GAS infection.

**Figure 3.**
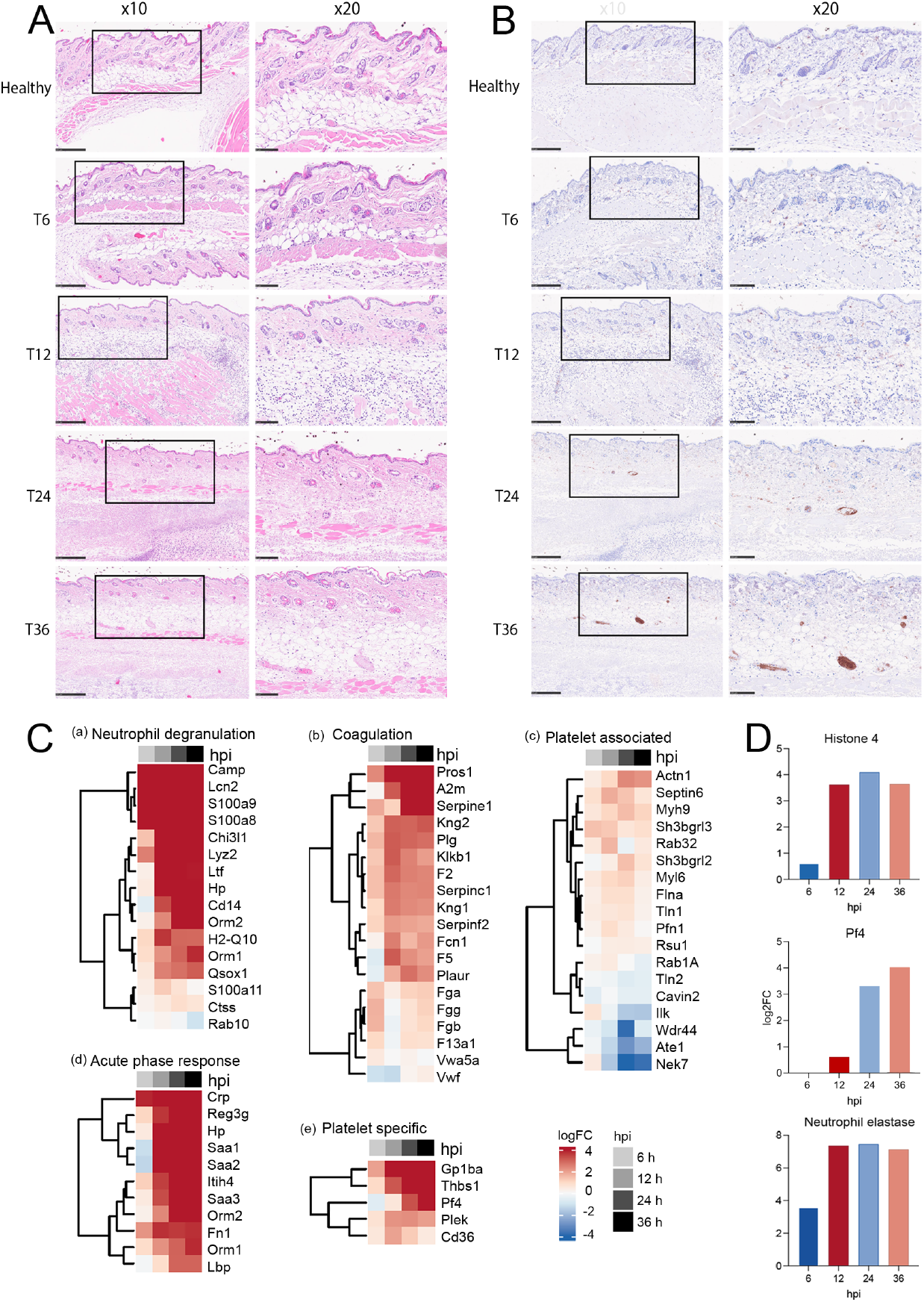
Platelets are enriched in skin biopsies during the late phase of *S. pyogenes-*induced skin pathology. Immunohistochemistry was combined with quantitative proteomics to assess the local inflammatory response and tissue pathology in skin biopsies. (A) Representative images illustrate the qualitative pathological changes over time in H&E-stained skin biopsies [Healthy (T0), T6, T12, T24, and T36] at two different magnifications. (B) Representative images illustrate immunohistochemistry of platelet accumulation (brown = CD41^+^ platelets) in the infected tissue over time. (C) Heatmaps depict log2 fold change for differential abundance of proteins in skin biopsies over time (T6-T36, grey scale above the heat maps) as compared to healthy mice (T0). Log2 fold change is plotted with upregulated proteins in red and downregulated in blue. Functionally enriched groups of proteins and showcase proteins from each group were defined based on Metascape analysis followed by manual curation of the generated datasets. (D) Temporal changes in marker proteins associated with neutrophil and platelet accumulation and activation in infected tissue.

### Trafficking of platelets to the liver during infection and sepsis

Immunohistochemistry and functional enrichment analysis revealed platelet accumulation in the skin at the later stages of infection. Platelet accumulation in distant organs was therefore investigated using immunohistochemistry and proteomics. Platelets were observed in the liver vessels at all time points but infiltrated into the tissue at 24 and 36 hpi (Fig. 4A). Some accumulation of platelets into the tissue occurred during late-stage organ damage; however, to a lesser degree than was observed in skin biopsies (Fig. 3B). Furthermore, platelet thrombi were not observed in the vessels of the liver as was previously observed for fibrin clots (Fig. 2Db). Comparison of the organ-specific proteome signature revealed that proteins involved in coagulation and fibrinolysis were enriched in liver and plasma at 24 and 36 hpi (Fig. 4B). Platelet-specific proteins were predominantly enriched in liver and plasma samples at the same time point (Fig. 4C), which supports the role of platelets in the coagulation system. The leukocyte pool is not expected to contain platelets, as these cells are removed in the washing procedures used for leukocyte isolation. Platelet proteins were not increased in the kidney samples; however, coagulation and fibrinolysis proteins were enriched, which implies that platelets do not accumulate in thrombi in kidney damage in this model. In the liver, platelet proteins peaked at 24 hpi but decreased again at 36 hpi. This agrees with the pattern of platelet activation reported using ELISA to detect CD62P in plasma (Fig. 1E), where plasma CD62P was highest at 24 hpi and decreased somewhat at 36 hpi. Distinct organ-specific patterns were observed for platelet proteins in plasma, liver, and skin. Pf4 (platelet factor 4), a chemokine released from activated platelets, was increased in liver and skin but not in plasma. Selp (CD62P) increased in plasma and liver at 24 and 36 hpi (Fig. 1). An increased abundance of Selp (CD62P) was not observed in skin samples (Fig. 3Ce); however, GP1ba was the most abundant platelet-specific protein observed. To investigate a potential mechanism underlying platelet accumulation, mouse whole blood was incubated with *S. pyogenes in vitro* in the presence or absence of platelet inhibitors targeting GPIbα (anti-GPIbα), GPIIb/IIIa (anti-GPIIb/IIIa or Integrilin), and P2Y_12_ (ticagrelor), and platelet-bacteria interactions were quantified by flow cytometry. Blockade of platelet GPIbα profoundly decreased bacterial binding to platelets, whereas anti-GPIIb/IIIa and P2Y_12_ inhibition decreased the interaction to a lesser extent (Supplementary Fig. 1). These findings suggest that GPIbα is a major contributor to platelet*-S. pyogenes* interactions. Collectively, these findings demonstrate organ-specific platelet accumulation and remodeling of platelet-associated proteomic signatures during invasive *S. pyogenes* infection, with GPIbα contributing to platelet-*S. pyogenes* interactions.

**Figure 4.**
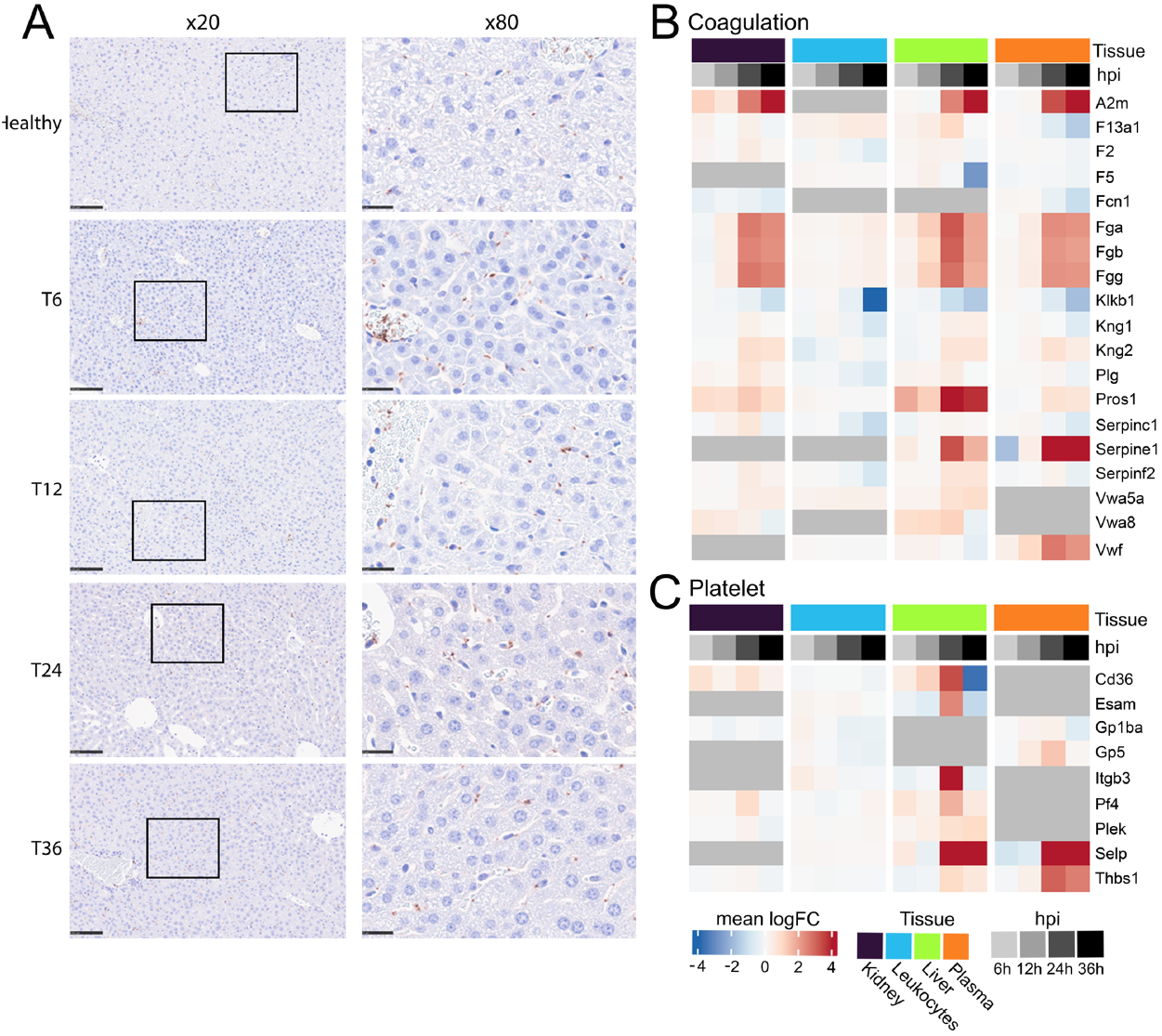
Temporal changes in platelet accumulation in infected and distant organs during sepsis. Immunohistochemistry was combined with quantitative proteomics to assess the role of platelets in the systemic inflammatory response and tissue pathology of sepsis-induced organ dysfunction. (A) Platelet accumulation in the liver as a representative of a distant organ was assessed using immunohistochemistry (brown = CD41^+^platelets) of organ biopsies (n=3) from mice at 0, 6, 12, 24 and 36 hpi. (B) Heatmaps depict DAPs associated with platelets in liver, plasma, leukocytes, and kidney samples over time (T6-T36 hpi, grey scale) as compared to healthy mice (T0). Log2-fold change is plotted for all DAPs with the direction of change colored in red (up) and blue (down). Functionally enriched groups of coagulation-(B) and platelet-(C) associated proteins and representative proteins from each group were defined using Metascape analysis followed by manual curation of the generated dataset.

### Intervention with antibiotics or antiplatelet therapy

To assess the effects of therapeutic intervention on the pathophysiological responses established in the model, mice were treated with antibacterial (meropenem) or an antiplatelet (ticagrelor) intervention at 12 hpi (Fig. 5A). This time point was chosen as the transition point between local and systemic infection when bacteria appear in distant organs, leukocytosis has occurred (Fig. 1), but organ damage has not manifested (Fig. 2). Meropenem is a broad-spectrum antibiotic that has previously been described as a therapeutic intervention in a mouse model of sepsis ^24^. Ticagrelor is a non-competitive, direct-acting platelet P2Y_12_-receptor antagonist that inhibits one pathway of platelet activation but does not cause a total blockade of platelet function, thus avoiding potential confounding effects of bleeding in the animals. Ticagrelor binds reversibly to the target to inhibit receptor signalling and subsequent platelet activation and does not require metabolism for activity. However, the parent compound is extensively metabolized, prolonging the efficacy and half-life of the drug. The dose of ticagrelor administered was determined to decrease murine platelet activation in a pilot experiment (Supplementary Fig. 2). The global proteome changes in liver, plasma, and skin showed a clear separation between naïve (uninfected) and infected/untreated mice (Fig. 5B). Mice administered with gold-standard antibiotic treatment with meropenem formed distinct clusters from infected/untreated mice in all organs investigated, demonstrating a clear treatment effect in the proteome (Fig. 5B). In contrast, antiplatelet intervention with ticagrelor resulted in a distinct proteome profile only in the liver, whereas plasma and skin proteomes largely overlapped with infected/untreated mice, which implies that antiplatelet therapy particularly impacted on the liver.

**Figure 5.**
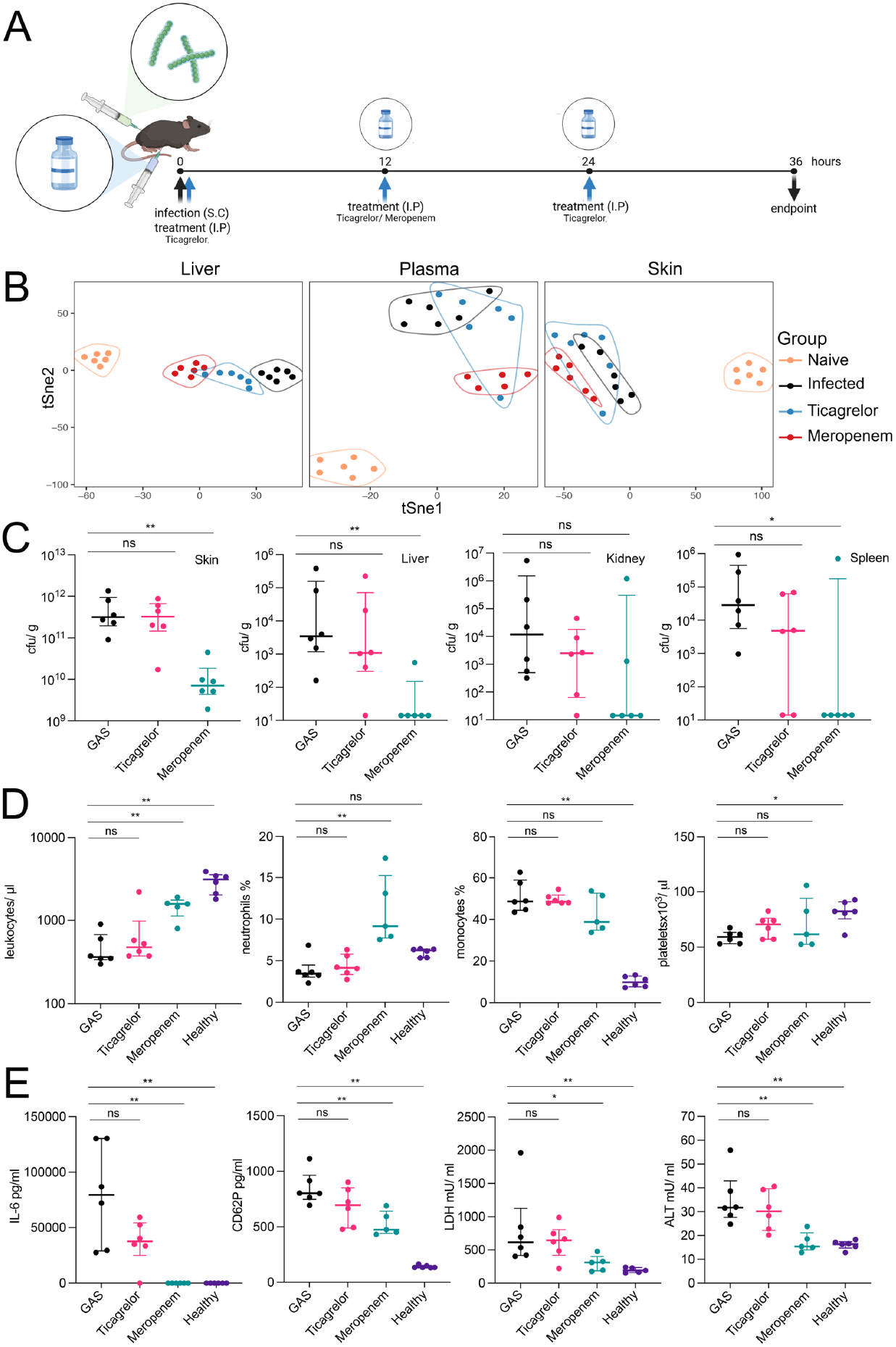
Response profiling in mice treated with antibacterial as compared to antiplatelet intervention. (A) Nine-week-old female C57BL/6J mice (n=6 per group) were infected subcutaneously on the flank with *S. pyogenes* AP1 or PBS (naïve) and, at 12 hours post-infection (hpi), treated with PBS (infected), antibacterial (meropenem), or antiplatelet (ticagrelor). Due to a short half-life of tecagrelor, ticagrelor-treated mice received an additional treatment at 24 hpi. All animals were sacrificed at 36 hpi and blood and organ samples were harvested. (B) tSNE plots from the full proteome responses, where samples were colored according to time points. (C) Bacterial load (cfu/g) was determined by serial dilution and viable count determination. (D) Blood cell distribution was analyzed by flow cytometry of total leukocytes (cells/µl), relative neutrophils and monocytes (percent (%) in the leukocyte gate, and total platelet count (cells/µl). (E) Markers of inflammation and organ damage were determined in plasma using ELISA; cytokine IL-6, platelet- and endothelium-derived CD62P, lactate dehydrogenase (LDH), and alanine transaminase activity (ALT).

Meropenem treatment significantly reduced bacterial burden at the local site (skin) and bacterial dissemination to the liver, kidney, and spleen (Fig. 5C). This also resulted in a significant improvement in leukopenia and neutropenia (Fig. 5D), together with diminished cytokine levels and platelet activation (Fig. 5E). Sepsis-induced thrombocytopenia was, however, not significantly improved by meropenem treatment (Fig. 5D). Importantly, sepsis-induced organ damage was reverted upon intervention with meropenem at 12 hpi (Fig. 5E). None of the parameters investigated were significantly affected upon intervention with antiplatelet therapy, indicating that the P2Y_12_ antagonist, ticagrelor, neither exacerbates nor significantly improves the progression to sepsis in this model. Interestingly, antiplatelet therapy did show a tendency toward improved thrombocytopenia, decreased cytokine levels, and decreased platelet activation in comparison to untreated animals (Fig. 5E), indicating a potential immunomodulatory role of platelets.

To quantify the impact of treatments on a proteome-wide level, we used a recently developed pharmacoproteomic intervention response score ^24^. The intervention response score was quantified as described in Supplementary Figure 3 by plotting the sepsis response against the treatment response signal to obtain slope and R^2^ as measures for treatment impact (Fig. 6A). The pharmacoproteomic intervention response was consistent with the cellular immunoassays (Fig. 5) that gold-standard treatment with antibiotics has protective effects. A reversion of the sepsis score was observed in liver and plasma in the meropenem-treated animals. A slope and R^2^ value of approximately 0.4 and 0.6, respectively, were found in liver and 0.3 and 0.5, respectively, in plasma (Fig. 6B), where a R^2^ value and a slope of 1 represent a full treatment response back to the uninfected state. Ticagrelor did not revert the sepsis response in plasma but had an impact on the liver response (Fig. 6A) with a slope and R^2^ value of approximately 0.2 and 0.2, respectively (Fig. 6B). Neither treatment impacted the sepsis score of skin significantly (Fig. 6A and B). The significantly reverted proteins in the liver across individual mice separated into three clusters (Fig. 6C). Clusters 2 and 3 were comprised of inflammatory and catabolic proteins upregulated in infected animals and meropenem reduced this effect while ticagrelor had little impact. Cluster 1 was comprised of proteins that were repressed during infection, and both treatments could diminish this effect. Metascape analyses revealed that proteins in Clusters 2 and 3 were mainly associated with acute-phase response, inflammation, and leukocyte function (Fig. 6D), while proteins associated with platelet activation and cellular homeostasis were enriched in Cluster 1 (Fig. 6D). Both treatments significantly reverted the total abundance of the proteins in Cluster 1 although with a more pronounced effect in the meropenem-treated animals (Fig. 6E). In the liver, both treatments partially reverted the abundance of proteins associated with cellular stress responses (Retreg2, Map2k3), as well as the immune mediator Ifit2 (Fig. 6F). Collectively, these findings demonstrate that meropenem broadly reverses sepsis-associated proteome changes, whereas ticagrelor exerts a more limited, liver-specific effect on the proteome.

**Figure 6.**
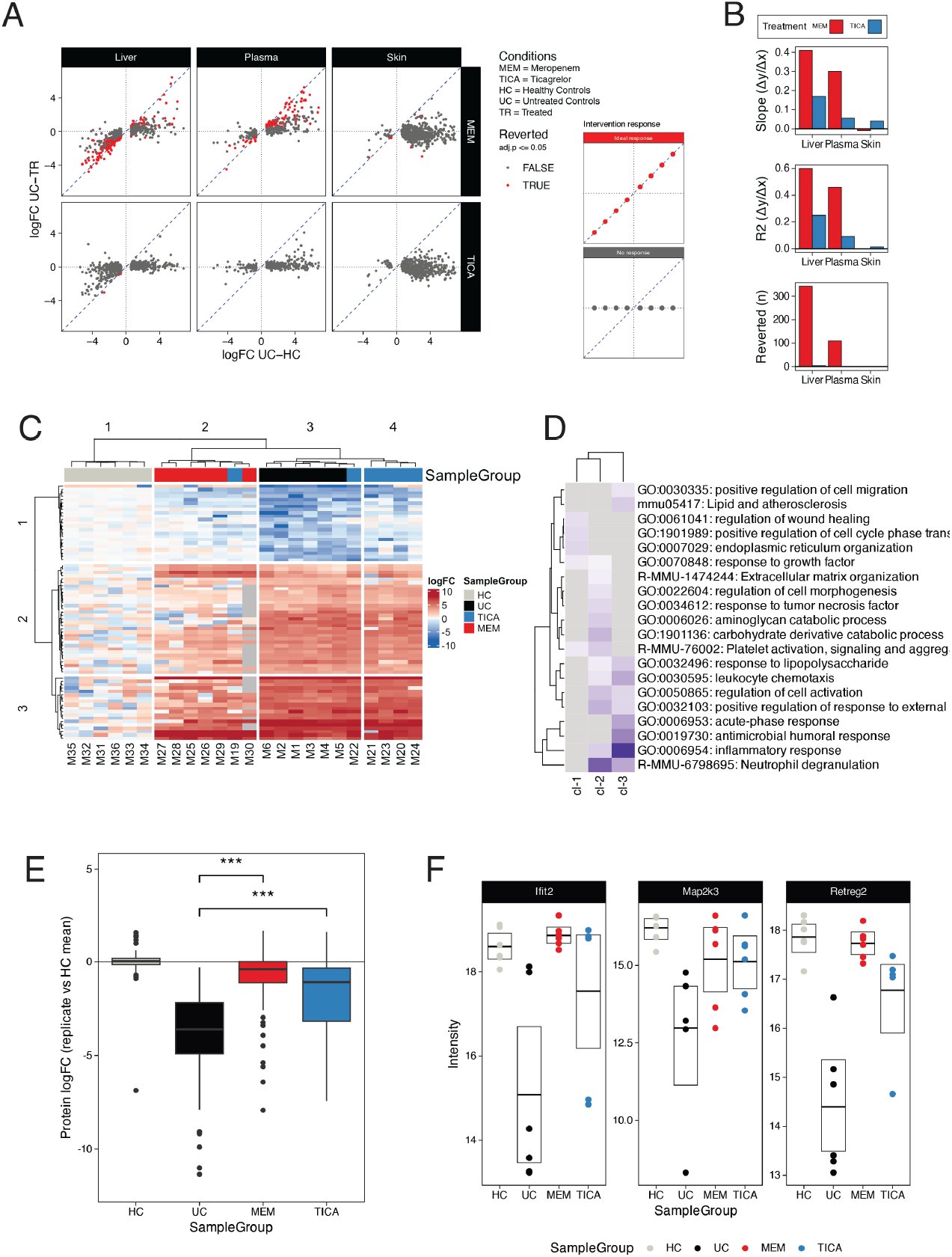
Pharmacoproteomic treatment response scoring. (A) Treatment response scatter plots comparing sepsis-induced protein changes (logFC UC-HC, x-axis) to treatment-induced reversion (logFC UC-TR, y-axis) for meropenem (MEM) and ticagrelor (TICA) across liver, plasma, and skin. Proteins significantly reverted toward healthy control levels are highlighted (red, adj.p ≤ 0.05). Dashed lines indicate no response (horizontal) and ideal response (diagonal). (B) Treatment response metrics by tissue: regression slope (Δy/Δx), coefficient of determination (R²), and number of significantly reverted proteins. (C) Heatmap of significantly reverted proteins across individual mice (columns) clustered into three protein groups (rows). Column annotation indicates sample group (HC, UC, TICA, MEM). (D) Pathway enrichment analysis (Metascape) of the three protein clusters from (C). (E) Distribution of protein log2 fold-change values for Cluster 1 proteins from (C) across sample groups. Pairwise comparisons were performed using the Wilcoxon rank-sum test. *** p < 0.001. (F) Protein abundance of representative Cluster 1 proteins (Ifit2, Map2k3, Retreg2) across sample groups.

## Discussion

*S. pyogenes* is an important cause of skin infection, and we have previously developed a mouse model of skin infection that progresses to sepsis ^25^. We have now used this model to map the pathophysiological organotypic changes during the progression from local infection to sepsis with a particular focus on platelets. We demonstrated that platelets are present at the local site of *S. pyogenes* skin infection, where they accumulate in the local vessels during the progressive necrosis of the tissue. Platelet and endothelial activation preceded organ dysfunction and platelets accumulated in the liver tissue as sepsis-induced organ damage progressed. Antibiotic treatment diminished the bacterial load and reduced organ damage, including a reversion of platelet activation and accumulation in the liver.

Antiplatelet treatment did not exacerbate sepsis, suggesting that the platelet activation inhibition is a viable target for modulation of sepsis without risking detrimental effects.

Our combined tissue, cellular, and molecular profiling of the mouse model revealed an early and late phase of disease that mirrors the pathophysiology of human sepsis. The first phase involved rapid recruitment of blood cells and acute-phase proteins to the local site of infection, mirrored by systemic changes in the blood cell distribution, leukocyte and liver proteome rearrangement, and systemic cytokine levels. This was followed by a second phase characterized by increasing necrosis in the skin, systemic tissue leakage, and cellular exhaustion represented by major downregulation of proteins in the liver and leukocytes, and pathological findings of organ damage in kidneys and liver that coincided with an increase in biomarkers of organ damage in plasma. Organ damage was particularly associated with an increase in proteins involved in innate immune functions and coagulation and this was mirrored at the cellular level as an accumulation of leukocytes, fibrin clots, and platelets in the liver. We demonstrated an increase in circulating platelet activation at the later stages of infection, which coincided with accumulation of platelets in the organs. This likely contributes to thrombocytopenia which is a hallmark of severe sepsis by removing platelets from the circulation. Large platelet aggregates accumulated in vessels at the local site of infection (skin); however, platelets were not associated with the fibrin-rich clots that accumulated in the vessels of the liver at late stages or sepsis-induced organ damage, suggesting that platelets are unlikely to be the primary drivers of coagulation-associated organ dysfunction in our model.

The rapid increase in platelet proteins in skin early in infection mirrors the increase in the classical immune cells, neutrophils. Immunohistochemistry demonstrated that platelets did not infiltrate into tissue to the same extent as neutrophils and were not colocalized with neutrophils in the tissue. Platelets were retained within the vessels of skin and, at later stages of disease, platelet aggregates were observed, which may contribute to hypoxia and necrosis in the tissue. Platelets have been reported to contribute to bacterial entrapment, release of pro-inflammatory mediators, and platelet-neutrophil interactions in the circulation ^13^. This potential interaction is supported by our findings that the platelet-derived neutrophil chemokine, PF4, and CD62P, an essential bridging molecule for the formation of platelet-neutrophil complexes, are elevated in our model. In particular, platelets stimulate formation of neutrophil extracellular traps (NETs) which contributes to bacterial containment and clearance ^26,27^. We observed increasing neutrophil numbers in the circulation preceding a pronounced neutropenia, corresponding with increased neutrophil extravasation into tissues and subsequent neutrophil degranulation, as observed in our proteomic profiling of skin, liver, and kidney. Neutrophil accumulation and NET formation in the organs will, however, also contribute to coagulation dysfunction and organ damage in the liver ^28^.

Our ex *vivo* experiments confirmed a specific interaction of *S. pyogenes* bacteria to platelets in blood, in a mechanism dependent on platelet GPIbα. Platelets have been shown to interact with different pathogenic microorganisms through a plethora of platelet expressed surface receptors resulting in platelet activation, degranulation, and aggregation ^29–32^. Thrombus formation in response to inflammation and infection has been proposed as an antimicrobial host response to entrap invading pathogens and prevent further dissemination ^33,34^. Platelets bind to *Listeria monocytogenes* bacteria in the circulation and transport them to the spleen for dendritic cell-mediated immune activation and clearance ^35^. Platelets also facilitate *Staphylococcus aureus* recognition and entrapment in liver sinusoids during systemic infection ^36^. However, as shown for infections with *Salmonella typhimurium*, immunothrombosis does not necessarily contribute to bacterial containment, instead prompting the formation of microthrombi and ultimately contributing to organ damage ^37^. Furthermore, virulence factors from distinct pathogens, including *S. pyogenes* may facilitate escape from fibrin or platelet thrombi ^38,39^. Hence, the interactions between platelets and GAS bacteria reported herein are likely to be species-specific and may contribute to the virulence of the pathogen.

Treatment with the P2Y_12_ receptor antagonist, ticagrelor, has previously been reported to result in lower levels of pro-inflammatory cytokines, reduced platelet activation, and prevented sepsis-induced kidney injury in a mouse model of sepsis ^17,18^. In our model, ticagrelor did not significantly impact the local response to infection or sepsis-induced organ damage. However, this antiplatelet therapy did partially revert the disease-associated increase in proteins associated with inflammation and cellular stress-induced liver dysfunction. Antibiotic treatment had a significant impact on both local infection and sepsis-induced organ damage by decreasing the bacterial burden and organotypic sepsis score including platelet enrichment in the liver at late stages of organ dysfunction.The high interindividual variability seen for the ticagrelor-treated group compared to the non-treated animals could in part be explained by the impaired microcirculation and associated organ damage, which could affect the pharmacokinetics of the administered treatment.

Altogether, we have determined the multifactorial pathways that contribute to the progression from local skin infection to dysregulated host immune responses, coagulopathy, and organ dysfunction in the kidney and liver. Platelet activation occurred late in disease progression and likely contributes to both sepsis-associated coagulopathy and immune dysregulation, thereby promoting collateral tissue damage and liver dysfunction.

## Acknowledgements

Gisela Hovold and Sara Wettemark are acknowledged for excellent technical assistance with the mouse model.

