## Supplemental Figures 1-4 for "Temporal changes in platelet activation and organ dysfunction during invasive *Streptococcus pyogenes* infection"

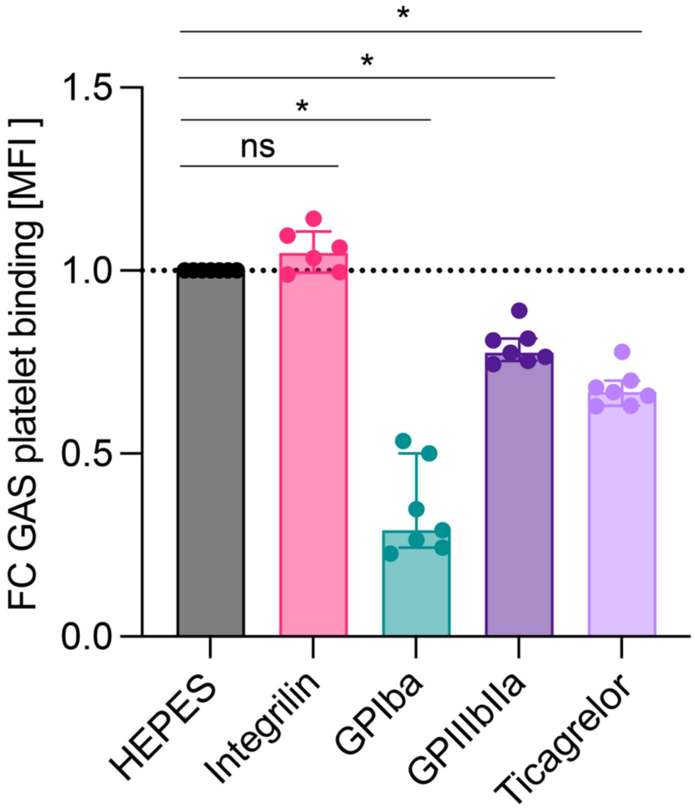


**Supplemental Fig 1**: The direct interaction between *S. pyogenes* and platelets was assessed in-vitro using flow cytometry of blood from healthy female C57BL/6J mice (n=7) stimulated with *S. pyogenes*. Data are shown as relative platelet-bacteria binding, measured by MFI of FITC labelled bacteria in the CD41 APC platelet gate. Pretreatment of blood with platelet receptor antagonists were used to determine the platelet receptors involved in bacteria binding; Integrilin (GPIIbIIIa blockade), anti-GPIba, anti-GPIIbIIIa antibodies and Ticagrelor ($P2Y_{12}$ blockade). Statistics calculated using Wilcoxon test. P values < 0.05 were considered statistically significant.

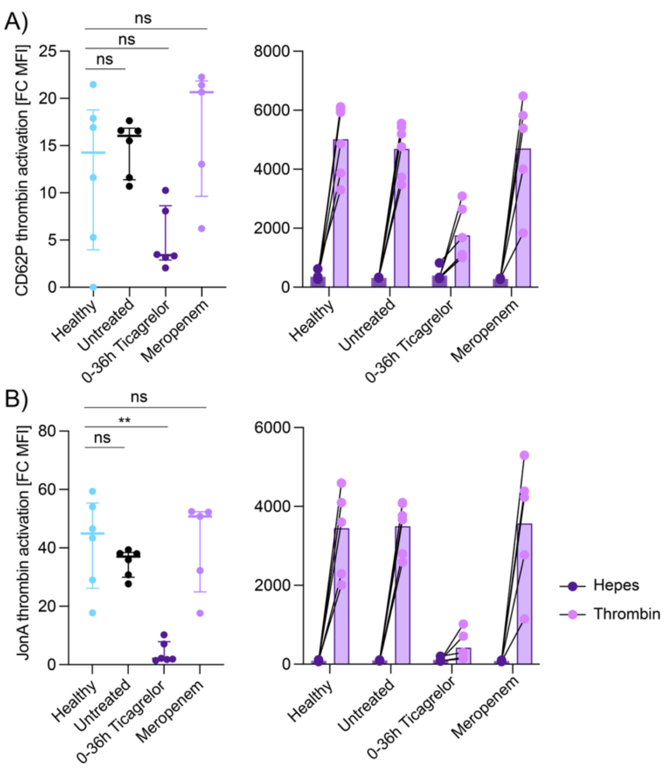


**Supplemental Fig 2**: Flow cytometry of blood from healthy female C57BL/6J mice (n=5) treated with tricagrelor or meropenum. Platelet activation was determined as thrombin induced upregulation of CD62P (A) and integrin activation (JONA) (B)

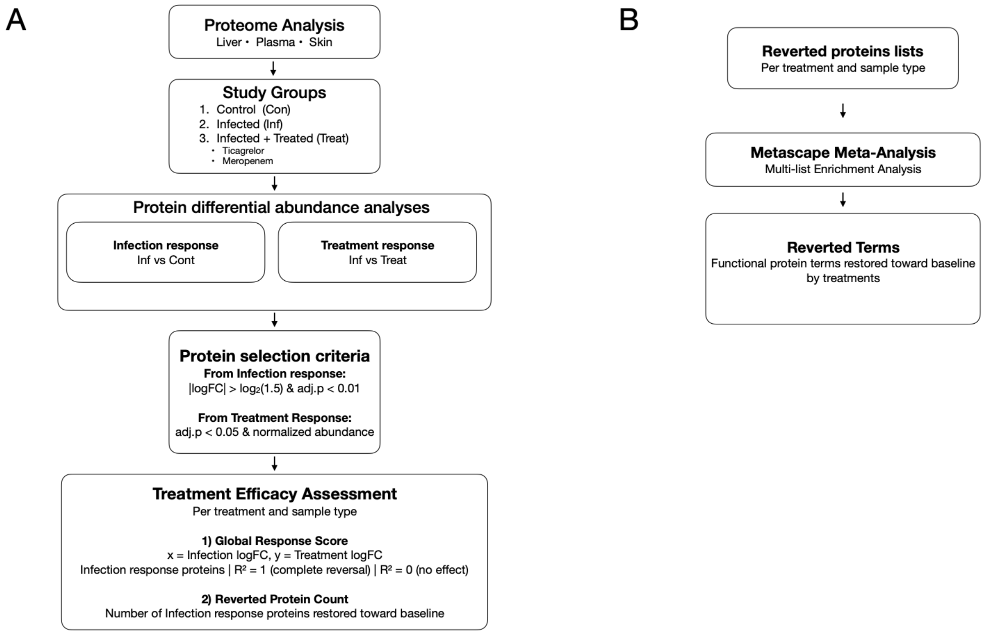


**Supplementary Figure 3**: Schematic overview of the generation of the sepsis intervention score

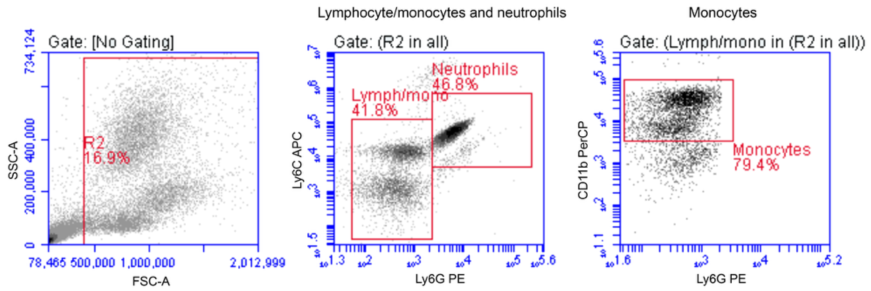


**Supplemental Figure 4. Representative gating for whole blood flow cytometry.**

Total leukocytes were gated (R2 gate) according to characteristic forward and side scatter (a). From total leukocytes (R2 gate), neutrophils were gated as Ly-6G and Ly-6C high (b). Lymphocytes and monocytes were gated together as Ly-6G low and Ly-6C intermediate/high. Monocytes were distinguished from lymphocytes with CD11b (c).
